# *Pseudomonas* superinfection drives Pf phage transmission within airway infections in patients with cystic fibrosis

**DOI:** 10.1101/2025.01.14.632786

**Authors:** Julie D. Pourtois, Naomi L. Haddock, Aditi Gupta, Arya Khosravi, Hunter Martinez, Amelia K. Schmidt, Prema S. Prakash, Ronit Jain, Piper Fleming, Tony H. Chang, Carlos Milla, Patrick R. Secor, Giulio A. De Leo, Paul L. Bollyky, Elizabeth B. Burgener

**Author notes:** Co-senior authors (Paul L. Bollyky, Elizabeth B. Burgener).

## Abstract

Pf bacteriophages, lysogenic viruses that infect *Pseudomonas aeruginosa (Pa),* are implicated in the pathogenesis of chronic *Pa* infections; phage-infected (Pf+) strains are known to predominate in people with cystic fibrosis (pwCF) who are older and have more severe disease. However, the transmission patterns of Pf underlying the progressive dominance of Pf+ strains are unclear. In particular, it is unknown whether phage transmission commonly occurs horizontally between bacteria within the airway via viral particles or if Pf+ bacteria are mostly acquired via new *Pseudomonas* infections. Here, we have studied *Pa* genomic sequences from 3 patient cohorts totaling 663 clinical isolates from 105 pwCF. We identify Pf+ isolates and analyze transmission patterns of Pf within patients between genetically similar groups of bacteria called “clone types”. We find that Pf is predominantly passed down vertically within *Pa* lineages and rarely via horizontal transfer between clone types within the airway. Conversely, we find extensive evidence of *Pa* superinfection by a new, genetically distinct *Pa* that is Pf+. Finally, we find that clinical isolates show reduced activity of the type IV pilus and reduced susceptibility to Pf *in vitro.* These results cast new light on the transmission of virulence-associated phages in the clinical setting.

## Introduction

Bacteriophages, viruses that parasitize bacteria, have complex relationships with their bacterial hosts. Purely lytic phages always lyse their bacterial hosts and can help clear bacterial infections (1,2). Conversely, lysogenic phages integrate their genetic material into the bacterial genome and reproduce via bacterial replication (i.e., lysogeny) or through opportunistic lysis (3,4). Finally, some phages can also produce new viral particles without killing the cell resulting in chronic infections of the bacterial cell (5–7). These phages can have a much more nuanced effect on bacterial populations, sometimes promoting treatment failure (4,8–10). One lysogenic phage that has been implicated in the pathogenic behavior of its bacterial host is the filamentous phage Pf, which is harbored by *Pseudomonas aeruginosa* (*Pa*) (9–11).

Pf phages are Inoviruses with a ssDNA genome encased in a filamentous structure a few nanometers wide and 1-2 micrometers long (5,6). They include Pf1 to Pf8 phages, in addition to other yet unassigned Pf phages, and are prevalent in multiple well-described lab strains of *Pa* (6,9,12,13). Filamentous phages like Pf are temperate and therefore have two possible routes of transmission. They have the ability to both integrate their 6000 to 15000bp genome into the genome of their bacterial host and to produce new viral particles that infect susceptible bacterial hosts by binding to the type IV pili (5,10,14). Filamentous phages form a non-lethal chronic infection during which progeny phages are produced and extruded without killing the bacterial host, instead of release by lysis as is the case with lytic phages (5,15).

The bacterial host of Pf phage, *Pa,* is a Gram-negative bacterium that is a common pathogen of humans. *Pa* is often resistant to multiple antibiotics, leading to its recognition as a pathogen of concern by the World Health Organization (16). It is present in the environment and is responsible for opportunistic infections of burns and diabetic wounds, for example, as well as infections in immuno-compromised individuals (17–19). In particular, *Pa* is one of most common bacteria found in lung infections in people with cystic fibrosis (pwCF) (20).

*Pa* is particularly problematic in cystic fibrosis (CF), a hereditary disease characterized by the disruption of ion channels (21). Symptoms include the accumulation of thickened mucus in the airways, leading to chronic bacterial infections and reduced lung function (22). Infection with *Pa* is a major predictor of morbidity and mortality in pwCF (23,24).

Worse outcomes in pwCF are associated with Pf presence (9,10,25), and phage-infected (Pf+) strains are known to predominate in older, more chronically ill pwCF. Similar findings have been reported in patients with chronic wounds infected with *Pa*, where Pf can be detected in the wounds, and more so in larger, slower to heal wounds (26). This dominance is thought to reflect selective advantages conferred by Pf through the modulation of phagocytosis (27–30) and the physical interaction of viral particles and biofilm polymers producing a crystalline organization of the biofilm, which reduces the efficacy of antibiotic treatment and thereby promotes antibiotic tolerance (31–34). Specifically, a charged-based interaction between negatively-charged Pf phages and positively-charged antibiotics results in the sequestration of these antibiotics away from bacteria and increased bacterial survival during antibiotic treatment (33,35). In addition, genes encoded by Pf result in modified quorum sensing signaling, inhibition of pyocyanin production and protection against superinfection by other Pf phages through suppression of the type IV pili (36–38).

Overall, Pf phages contribute to the fitness and pathogenicity of *Pa,* especially under antibiotic treatment, and favor the development of chronic infections (10,11,34). In pwCF, this manifests into Pf+ infections being associated with advanced disease and worse pulmonary exacerbations (9). There are also reports that Pf phage is associated with bacterial phenotypes linked to reduced virulence and chronic infection (39–41). Yet, little is known about the transmission of filamentous phages in the clinical setting, which involves heterogeneous environments (42–44) and multiple bacterial clone types—closely-related isolates—both within and between patients (45–50).

The lungs of patients with cystic fibrosis present a unique environment with bacteria and phages navigating a highly complex spatial structure (42–44,51). In addition, *Pa* has been shown to adapt to the CF airway and exhibit changes in metabolism, antibiotic resistance, motility and other virulence factors that could affect Pf infection (44,46,52). While the prevalence of Pf-infected bacteria increases with patients’ age (9), the mechanisms driving this increase are unclear. In particular, we do not know whether Pf+ strains are generally acquired directly from the environment and become more prevalent in a patient with time as a result of the competitive advantage of Pf+ bacteria, especially under antibiotic treatment, or whether horizontal transmission of phages among bacteria within a patient’s airway commonly contributes to this increase.

In this study, we focus on Pf phages of *Pa* infecting the airways of pwCF. We use bacterial genomes from three different patient cohorts to describe patterns of infection by Pf and investigate how Pf phages spread within patients. We use genomic data from bacterial samples from two different cohorts of pwCF already published in the literature (46,47) and newly-sequenced samples from a third cohort of patients from the Cystic Fibrosis Center at Stanford to investigate patterns in Pf transmission in *Pa* lung infections in patients with cystic fibrosis. We ask if Pf phages are often transmitted within patients or if Pf+ infections usually represent new clone types. We find some evidence for horizontal transmission of Pf in the phylogenies, but do not observe Pf transmission for more than 95% of clone types within patients. These results suggest that new Pf+ infections are typically caused by a new bacterial infection with a genetically distinct Pf+ *Pa* rather than horizontal transmission of Pf to an established *Pa* infection.

## Results

We collected and sequenced bacterial isolates from pwCF treated at the Cystic Fibrosis Center at Stanford (California cohort). In total, 163 bacterial samples were then sequenced, across 67 pwCF in nearly 3 years (Figure 1A). We combined these with publicly available sequences from a cohort of Danish and a cohort of Italian pwCF. These datasets contain 474 and 26 isolates from 34 and 4 pwCF, across 11 and 19 years respectively (Figure 1A) (see Marvig, Dolce, et al. 2015; Marvig, Sommer, et al. 2015). Isolates from each cohort cluster into groups of high genetic similarity, that have recently diverged within individuals and are characterized by fewer than 10,000 SNPs (Figure 1B). We refer to these groups as clone types. If a new clone type appears in a subject we consider that a superinfection.

**Figure 1.**
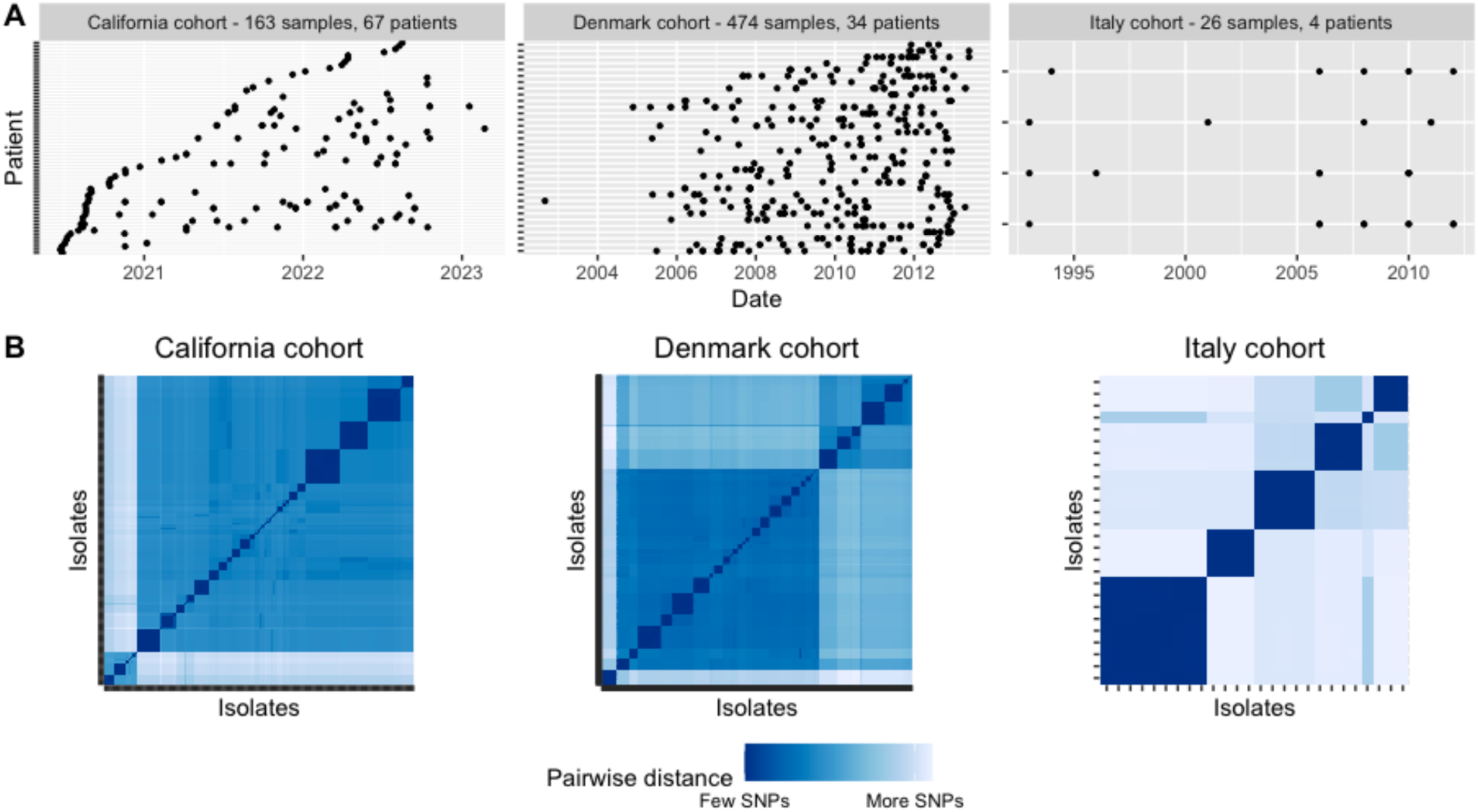
Overview of the three patient cohorts used in this study. **(A)** Time-series of sample collection from patients for each cohort. **(B)** Genetic distance matrix showing pairwise SNPs between each isolate for each cohort. Clinical isolates cluster into groups of high genetic similarity called clonal types. Genetic distance scale varies between cohort.

### Pf phages are found in high proportions across the three patient cohorts

We first asked how prevalent Pf phages are across patient cohorts in different countries. Lineages of Pf phages target distinct integration sites to integrate themselves into bacterial genomes (13). A single bacterium can thus be infected by multiple Pf phages (53).

We used the presence of five highly conserved Pf genes—*PA0718, PA0719, PA0720, PA0721* and *PA0727*—to detect Pf prophages in the chromosome of *Pa* clinical isolates (13). Using a threshold of 75% total coverage of these core genes, we found that between 65 and 69% of isolates from each *Pa* patient cohort were Pf+ (Figure 2A). These results were not sensitive to changes in coverage threshold between 35 and 85% (Figure S1).

**Figure 2.**
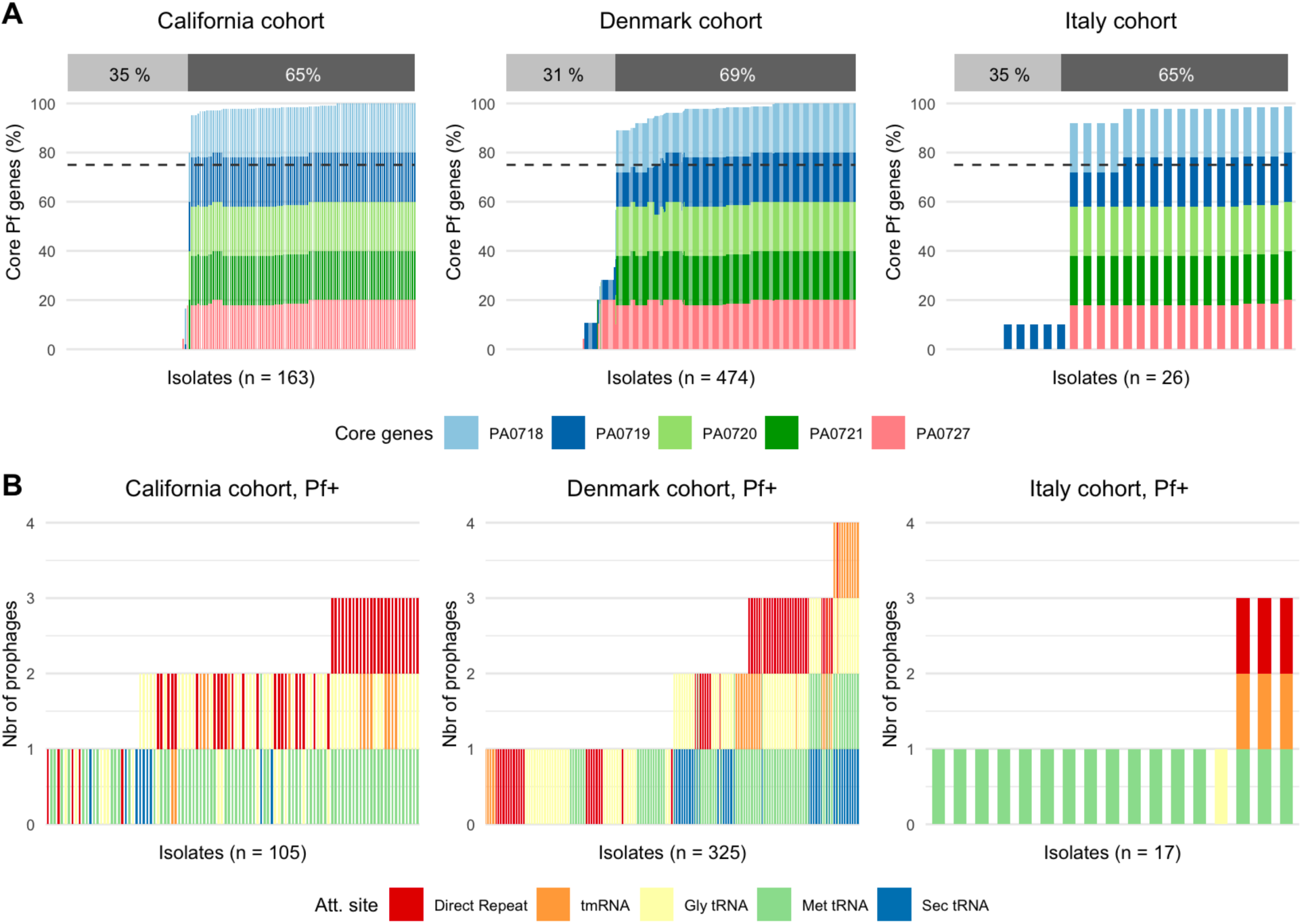
Majority of *Pa* infected by Pf phages. **(A)** Percent of core Pf genes PA0718, PA0719, PA0720, PA0721 and PA0727 found in isolates for each patient cohort. The percentage of Pf+ isolates, defined with a minimum average coverage of 75% for these five core genes, is shown above each graph. **(B)** Number of prophages found in Pf+ isolates, as defined above, and attachment site used for each patient cohort.

The number of Pf prophages infecting each isolate was then determined using variation in the Pf integrase gene *intF* (Figure 2B). Each integration site in the *Pa* chromosome can be targeted by different Pf phages with the corresponding integrase. We thus used the five different integrases described in the literature (see **Methods**) to categorize Pf phages into five different Pf ‘types’ and to count the number of phages infecting each isolate. We validated this approach using long-read sequencing of twelve isolates with different predicted number of phages and did not find any instances of multiple infections by phages with the same integration site in a single bacterial isolate. Coinfection occurred in many isolates, with up to 4 different Pf prophages per isolates in the cohort from Denmark (Figure 2B). The most common integration sites used by Pf phages were Met-tRNA and Gly-tRNA (32% and 28% respectively), followed by direct repeats (21%). These integration sites are used by well-described Pf phages, with reference Pf phages Pf4, Pf5, Pf6 and Pf7 using Gly-tRNA, direct repeats, Met-tRNA and Met-tRNA respectively (13). The least commonly found integration sites were tmRNA and Sec-tRNA at 10% and 9% respectively.

### Gain or loss of Pf phage by clone types is rare within patients but more prevalent outside of patients

We mapped the presence of Pf phages with different integrases onto phylogenetic trees of the bacterial isolates for each cohort. Pf+ and Pf- bacteria were present across the phylogenetic tree, while the number of Pf phages was usually conserved within clone types in patients (eg. clone types CA14, DK14 and IT01, Figure 3A,B,C). Specifically, 98%, 95%, and 100% of isolate pairs from the same clone type in the same patient were infected by the same number and same type of phages, as defined by their integrase, for the California, Denmark and Italy patient cohorts respectively (Figure 3D). Isolates of the same clone type in a patient were significantly more likely to be infected by the same Pf phages than other isolate pairs for all clinical cohorts (Chi-square test, p < 0.001 for all). Furthermore, Pf phages of the same type, as determined by their integrases, infecting bacteria of the same clone type in the same patient had a significantly lower number of non-identical nucleotides than for Pf phages infecting different clone types or different patients for both the California and Denmark cohorts (Wilcoxon rank-sum test, p < 0.001 for both, Figure 3E). In other words, Pf phages of the same type in the same patient were likely to have diverged within that patient, rather than to represent a new Pf phage from the same type from the environment or other patients. Together, these results suggest that clone types do not typically gain new phages, whether of the same type or not, at the timescale of divergence within a clone type over multiple years in patients. Note that the low number of mutations observed in Pf phages due to their small size (∼10,000bp) did not allow us to evaluate Pf transmission of a particular phage type within a bacterial clone type.

**Figure 3.**
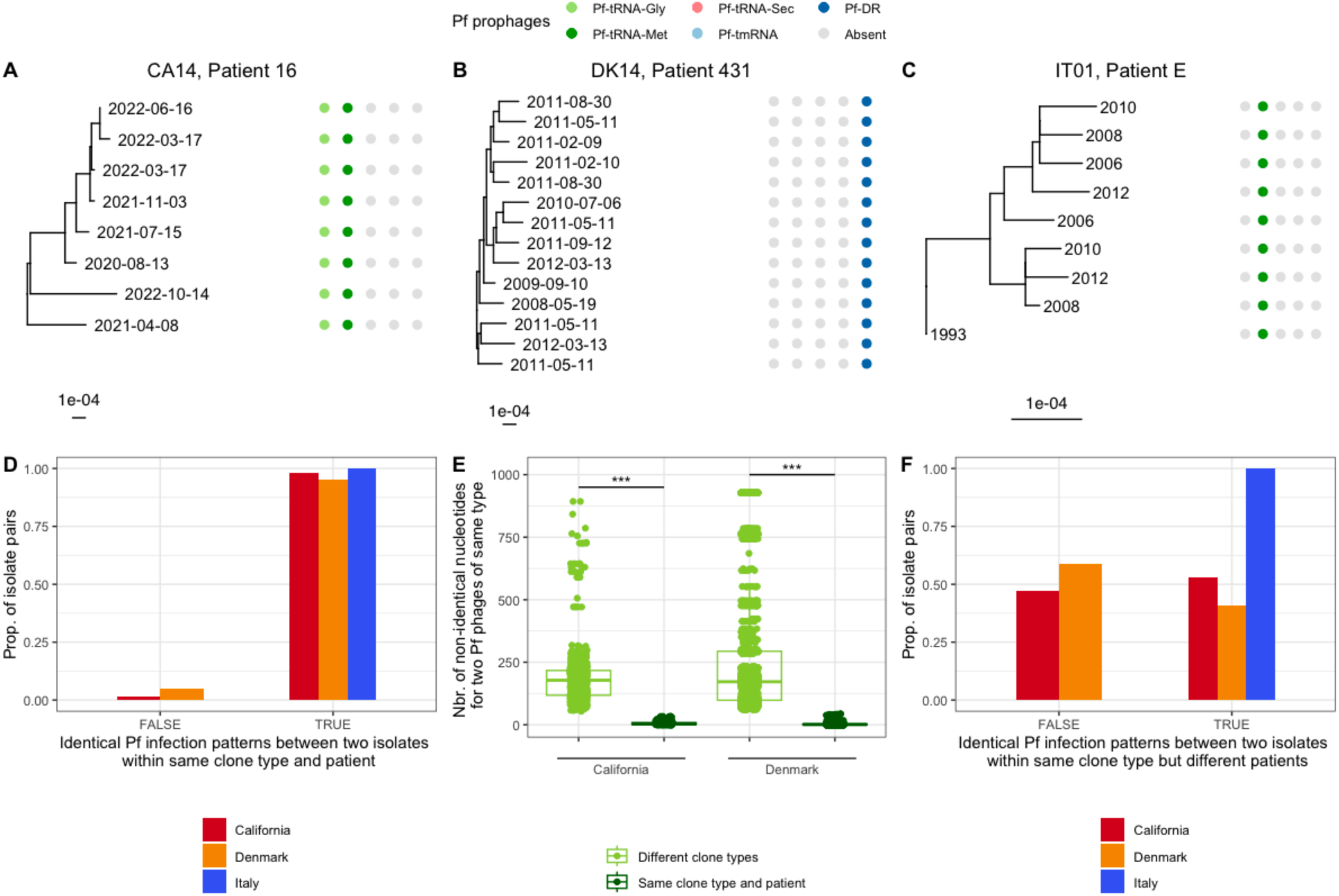
Pf prophages are maintained within clone types within patients. Presence or absence of five different Pf types, as defined by their integrase, for one clone type and patient in the California cohort **(A**), Denmark cohort **(B)** and Italy cohort **(C).** Sample dates are shown at the end of each branch. **(D)** Proportion of *Pa* isolate pairs of the same clone type and in the same patient with identical Pf infection patterns (same number and type of Pf phages, as determined by their integrase). **(E)** Number of non-identical nucleotides for pairwise comparisons of Pf genomes of the same phage type (using the same integrase) for Pf infecting the same patient and *Pa* clone type and for Pf infecting different clone types. **(F)** Proportion of *Pa* isolate pairs of the same clone type but in different patients with identical Pf infection patterns (same number and type of Pf phages, as determined by their integrase).

We then asked if Pf transmission was more common outside of patients. We observed that 53 %, 41% and 100% of isolates from the same clone type but different patients were infected by the same Pf phages in the California, Denmark and Italy cohorts respectively (Figure 3F). The proportion of isolates that had the same number and same type of phages was significantly lower for isolates of the same clone types but in different patients than in the same patients for the California and Denmark cohorts (Chi-square, p < 0.001 for both). There were no isolates of the same clone type in different patients for the Italy cohort. These data indicate that Pf transmission may be more common outside than inside of patients and may be prevented by phenotypes associated with the CF airway.

### Infection by a new Pa clone type, rather than horizontal transmission, is responsible for most new Pf+ infections in pwCF

Many patients carried either only Pf- or Pf+ isolates over the study period, with 60%, 50%, and 25% of patients with multiple samples seeing no change in the number of Pf prophages for the California, Denmark and Italy cohort respectively (Figure 4A). When patients were infected with isolates with different number of phages 64%, 72% and 100% of the changes in Pf prophage copy numbers were associated with a change in clone type for the California, Denmark and Italy cohort respectively (Figure 4B, 4C). All changes between Pf- to Pf+ (independently of prophage number) represented a change in clone type in all cohorts, except for patient 31 in the California cohort (Figure 4C). Pf+ infections in patients were thus usually caused by infection by a new clone type, rather than by the gain of a new Pf prophage by an existing Pf- clone type.

**Figure 4.**
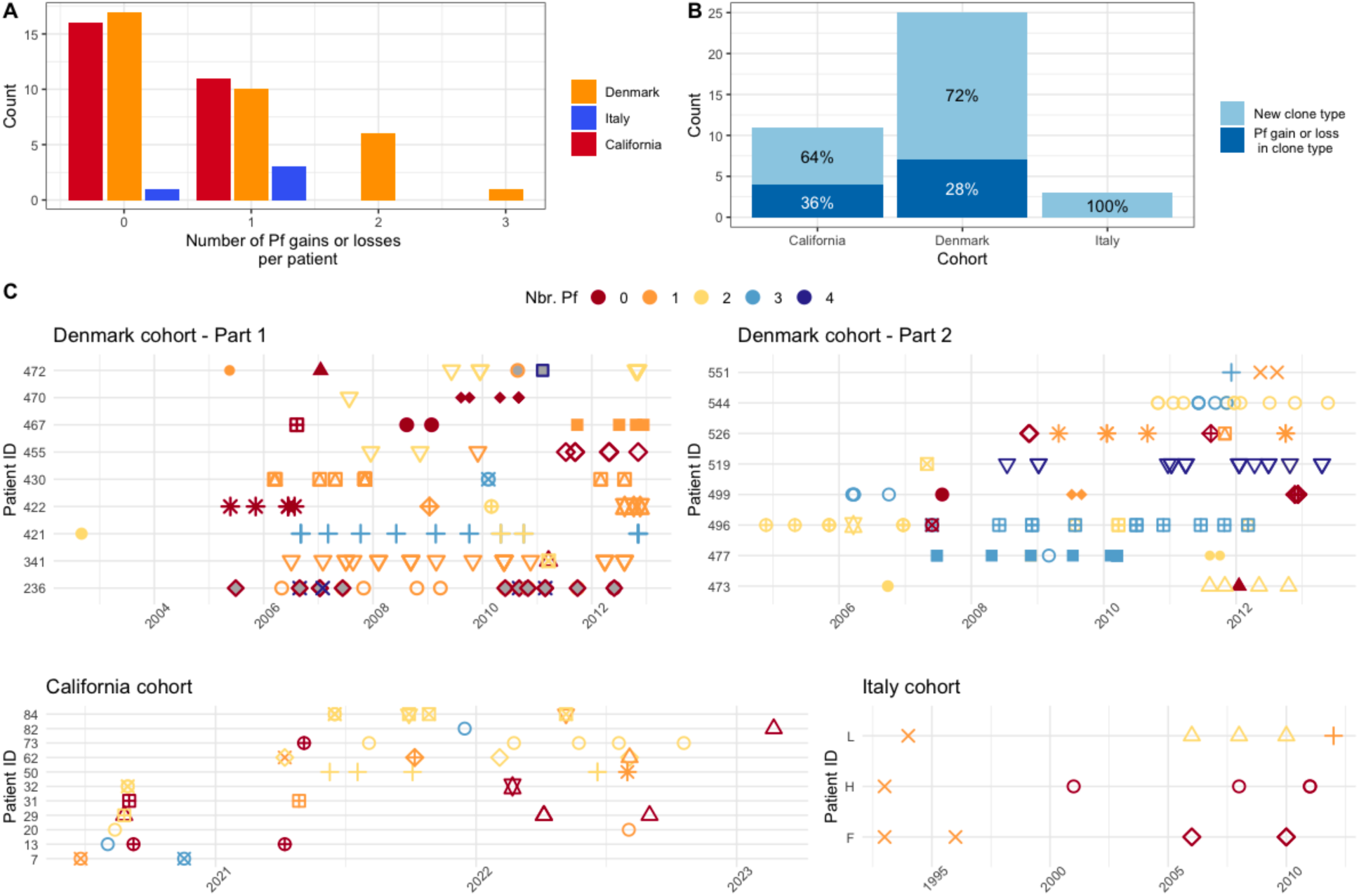
The majority of changes in the number of Pf phages in patients are associated with colonization or dominance of new clone types. **(A)** Number of changes in the number of Pf phages per patient. **(B)** Percentage of the changes in the number of Pf phages that are due to a change in clone type or to infection of a clone type by Pf (horizontal transmission). **(C)** Time series of the number of phages and clone types for patients with at least one change in the number of phages over time. Different shapes represent different clone types while color indicates the number of Pf prophages found in that isolate. The same shapes were used to represent different clone types between the left and right panel for the Denmark cohort and between the different cohorts.

Finally, we observed a dichotomy in clone type persistence between patients. Patients with changes in clone types and Pf status often went through many of these changes through time (Figure 4C), while other patients remained infected by the same clone type for decades (Figure 4A, Figure S2). Time series for patients without a change in Pf are available in Figure S2.

### Clinical isolates do not twitch and show decreased susceptibility to infection by Pf in vitro

Pf phages rely on type IV pili on the surface of bacteria to attach to and infect bacterial cells (14,36). These pili allow *Pa* to twitch and move along a surface (Figure 5A). We selected Pf- isolates (CPA0056, CPA0139 and CPA122) from patients infected with both Pf- and Pf+ clinical isolates over time and asked if they had maintained functional pili and could be infected *in vitro.* We found that these clinical isolates had reduced pili function, as indicated by a smaller twitching radius compared to the laboratory strain PAO1 (Figure 5B, p < 0.05 for all, Wilcoxon test). Finally, we investigated the susceptibility to Pf infection of clinical isolates compared to PA01. While Pf does not need to lyse its host during endogenous replication, lysis is commonly observed during Pf superinfection at high multiplicity of infection (MOI) (11,36,54). We observed that Pf causes plaques or reduced growth at 10^3^ PFU/ml or above for PAO1, while 10^11^ PFU/ml or above were required to inhibit growth or cause plaques for all clinical isolates tested (Figure 5C).

**Figure 5.**
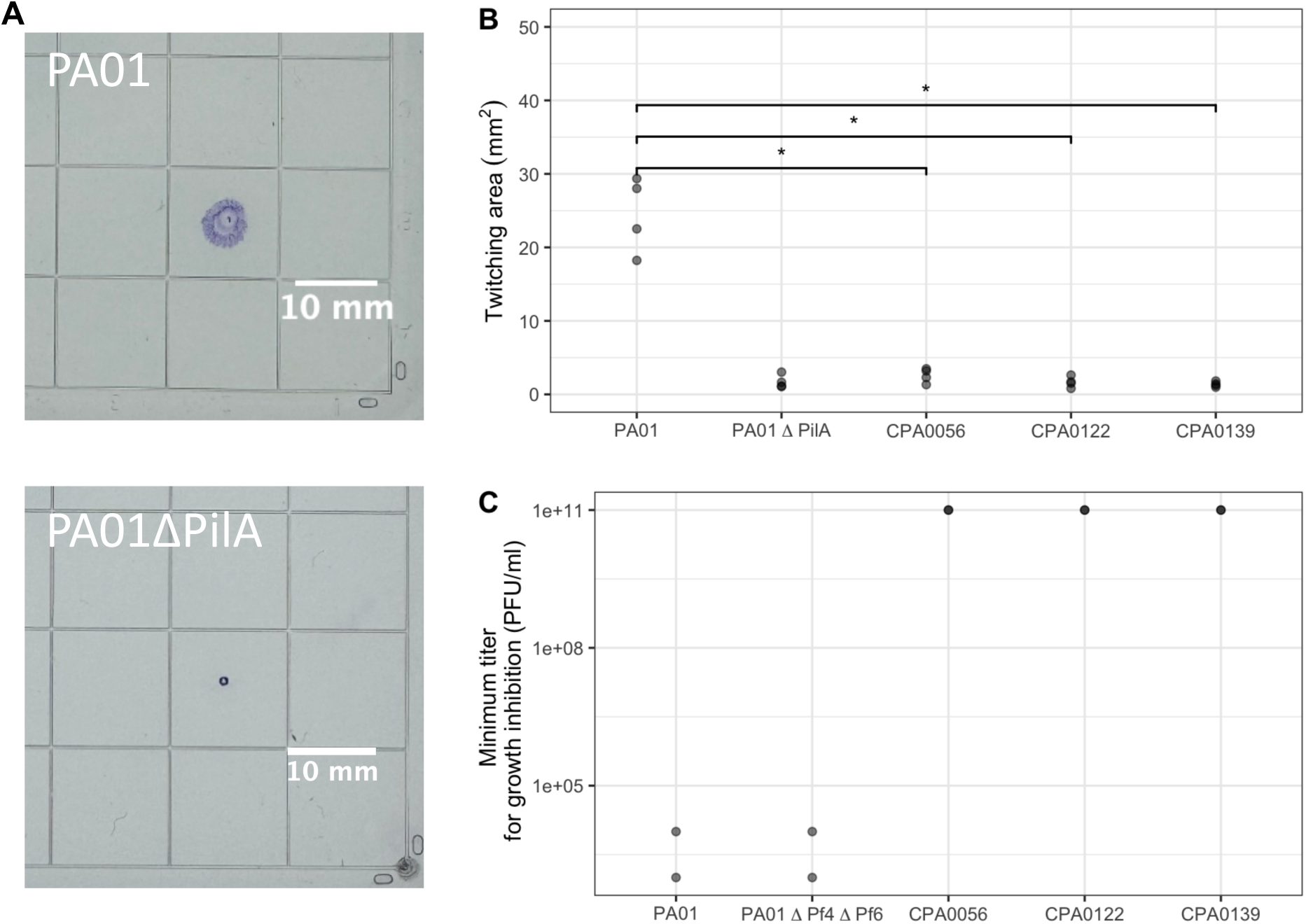
Clinical isolates do not twitch and are less susceptible to Pf than PA01. Clinical isolates were selected from patients that were infected by both Pf- and Pf+ isolates. **(A)** Positive (PA01) and negative (PA01ΔPilA) controls for twitching assays. **(B)** Twitching area for PA01, PA01ΔPilA and three Pf- clinical isolates. **(C)** Minimum titer at which plaques were present or *Pa* growth was inhibited for PA01, PA01ΔPf4 ΔPf6 and three Pf- clinical isolates.

## Discussion

Temperate phages are known to contribute to bacterial pathogenesis and to influence human health but their transmission dynamics can be complex. Here, we have investigated the transmission patterns of the temperate phage Pf in a clinical context, specifically in three independent cohorts of pwCF. We found that 60-70% of clinical isolates were infected with Pf and that most of these Pf+ isolates were infected with more than one Pf phage, as determined by the presence of different Pf integrases (13).

We find that Pf phage is typically maintained within *Pa* clone types. Within our dataset, the number and type of Pf prophages infecting each clone type was stable for more than 95% of *Pa* clone types within patients, sometimes over decades. Pf phages with the same integrase in a clone type were more genetically similar to each other than to other phages with the same integrase, suggesting that the transmission of Pf phages, even of the same type, does not commonly occur between *Pa* clone types within patients. In contrast to vertical transmission, horizontal transmission between different clone types is limited in pwCF. The majority of clone types did not gain Pf phages during chronic infections in patients, even when these lasted more than 10 years.

Any analysis of the transmission patterns of Pf phages within a clone type, however, was limited by the low rate of mutation of *P. aeruginosa* in patients, estimated at 2.6 SNPs/year (47). This is reflected in slow divergence at the bacterial level in pwCF and even slower divergence of phages. Phages within clone types often shared 100% of their DNA, making it impossible to track phage transmission within a clone type. In addition, the presence of well-conserved genes (eg. PA0721) close to genomic areas of high diversity (eg. coat proteins) in the Pf genome resulted in poor Pf genome reconstruction during genome alignment and limited our ability to establish transmission outside of patients. Our work established the need for methods tailored to Pf phages in order to fully capture the diversity of Pf phage infections and identify coinfections. Future work could combine short read and long read sequencing for optimal assembly and identification of prophages and investigate Pf transmission over larger scales.

Additionally, as these isolates were cultured in the clinical microbiology lab, typically 1-2 different appearing colonies were identified and reported and subsequently banked. Using sequencing, rather than visual appearance, it has been described that pwCF can harbor 10s-100s of clones within the individual (55,56). We may be missing some aspects of Pf phage transmission by not evaluating more clone types per subject, however we are likely evaluating the dominant clones within each patient.

Despite the low prevalence of Pf transmission between different clone types within patients, more than 50% of patients saw gains or losses of Pf phages over time. The majority of these changes were associated with a change in clone type, rather than Pf infection of a clone type previously detected in that patient. We investigated factors that could affect the absence of transmission of Pf in patients. Previous work shows that Pf is present in the sputum of some pwCF (9), indicating that Pf phages virions are actively produced by *Pa* in clinical infections. Lack of Pf virion production is thus unlikely to explain the near-complete absence of Pf transmission in patients. Pf has been shown to prevent superinfection by additional Pf phages by inhibiting the function of type IV pili, which are used by Pf as receptors (36). However, the majority of Pf- clone types also did not gain new Pf phages in patients, suggesting the presence of additional mechanisms preventing superinfection.

The CF airway is a complex environment and it is possible that the spatial separation between different clone types in the lung, arising either from airway structure or properties of the *Pa* biofilm, would extend to their phages. In addition, *Pa* has been found to undergo many adaptations in the CF airway, generally moving to a less virulent phenotype, including downregulation of pili expression and reduced motility (46,57–59), both of which could affect the ability of Pf to reach other bacteria and to attach to the type IV pilus to infect *Pa*. Here, we tested twitching ability, as a proxy for type IV pili function (36), and susceptibility to Pf of clinical isolates sampled from pwCF infected with both Pf+ and Pf- isolates. We found that all clinical isolates tested had reduced twitching ability and reduced susceptibility to Pf at clinically-relevant titers compared to the reference PAO1. These results suggest that the reduction in twitching often observed in isolates from established *Pa* infections may prevent superinfection by Pf phages.

Given the clinical associations of Pf phage with poor outcomes in CF (9), these data indicate that the initial acquisition of *Pa* with a Pf+ isolate may carry prognostic value, perhaps indicating a worse prognosis or trajectory if Pf phage is present. Additionally, the effect of Pf phage on biofilms and interaction with antibiotics (31,35) could also indicate high likelihood of establishing chronic infection, or low likelihood of successful eradication. Our groups are currently studying these hypotheses in longitudinal cohorts and mechanistic studies in the laboratory.

In summary, we have examined the transmission patterns of Pf phage, in cohorts of patients with CF and observe that new Pf+ infections are typically caused by new bacterial infections rather than horizontal transmission of Pf from a co-infecting clone type. Moreover, Pf+ and Pf- strains can coexist within patients and the balance of these strains within individuals can change over time. These results cast new light on the transmission of virulence-associated phages in pwCF.

## Methods

### Collection of P. aeruginosa isolates at the Cystic Fibrosis Center at Stanford (California Cohort)

From June 2020 to June 2023, *Pa* isolates from respiratory cultures from individuals with CF were identified and banked with patient consent for biobanking under IRB #11197. Sex was not considered as a biological variable.

### DNA extraction and sequencing

This study includes 163 new clinical isolates from 67 patients at the Cystic Fibrosis Center at Stanford. DNA was extracted using the DNA Easy Kit (Qiagen, 69504, Hilden, Germany) and sequenced on an Illumina NovaSeq (100bp paired-end) and an Illumina NextSeq (150bp paired-end) (60). We also extracted DNA from 12 samples using the Monarch HMW DNA Extraction Kit for validation using long-read sequencing. Long-read sequencing was performed using Nanopore R10.4.1 flow cells.

### Twitching assays

Twitch motility was assessed as previously reported (27). PAO1, PAO1ΔpilA, or indicated clinical isolates *P. aeruginosa* were stab inoculated through a 1.5% agar LB plate to the underlying plastic dish. After incubation for 24 hours, the agar was carefully removed, and the zone of motility on the plastic dish was visualized and measured after staining with 0.05% Coomassie brilliant blue. Twitching area was measured using ImageJ.

### Plaque assays

Plaque assays were performed using PaO1, PAO1ΔPf4ΔPf6 or indicated clinical isolates of *P. aeruginosa* as recipient strains on LB agar plates. These plates were prepared by adding 5ml of soft top agar media, consisting of tryptone (10 g/L), NaCl (10 g/L) and agar (5 g/L), mixed with 100ul of the *P*. *aeruginosa* recipient strain (OD600 0.4-0.6) per LB agar plate. Plaque assays were performed by serially diluting purified Pf4 in phosphate buffered saline (PBS) and spotting 10µl onto the prepared top agar plates. Plaque forming units were quantitated after 18-24 hours of growth at 37 °C.

### Sequence acquisition and assembly

Raw reads for isolates from the Italy patient cohort were downloaded from the Short Read Archives. Raw reads from the California and Italy patient cohorts were trimmed using trimmomatic (61) for the following parameters: java -jar /Trimmomatic-0.39/trimmomatic-0.39.jar PE -threads 8 -phred33 “$input1” “$input2” “$output1” “$output2” “$output3” “$output4” ILLUMINACLIPTrimmomatic-0.39/adapters/TruSeq3-PE.fa:2:30:10:2:True LEADING:3 TRAILING:3 MINLEN:36. We used trim_galore for paired reads to remove Nextera adapaters for raw reads from the Illumina NextSeq. Trimmed reads were then assembled with SPAdes using –isolate and --cov-cutoff auto (62). Assembled sequences were acquired directly from Marvig et al. for the patient cohort from Denmark.

### Clone type identification

Isolates were separated into genetically-similar clone types according to the methods described in Marvig et al (46). Briefly, we used the dnadiff command from mummer/4.0.0rc1 to obtain the number of mutations between each pair of isolates (63). Isolates were then assigned to an existing clone type if they had fewer than 10,000 SNPs when compared to other members of that clone type or assigned to a new clone type if they had more than 10,000 SNPs compared to all other isolates.

### Pf prophage identification

A custom database was built for the conserved Pf genes PA0718, PA0719, PA0720, PA0721 and PA0727 using the command makeblastdb from the blast package. We then used blastn to look for the presence of these five genes in all the bacterial isolates with options -word_size 28 -evalue 0.005 and -outfmt “6 qseqid sseqid pident length qstart qend sstart send sframe evalue qlen slen qseq”. We calculated the coverage (percent of gene length matched by blast) for each gene in R. Isolates were labelled as Pf-positive if the total coverage across the five genes averaged to more than 75%. Our results are not sensitive to changes to this threshold between 35 and 85% (Figure S1).

The number of Pf phages infecting each isolate was determined using the phage integrase gene PA0728, building on the assumption that an integration site can only be used by one phage at a time. We only recorded the presence of a phage if the integrase PA0728 was found next to the gene PA0727 (within 2000 bp), along with either the corresponding excisionase or integration sites. Phages were further classified as containing either the isoform A or B of the CoaB protein. The five different integrase, three different excisionase and two different CoaB proteins described by Fiedoruck et al. were identified using blastx, based on amino acid sequences (13). We looked for the presence of the five different integration sites using blastn with a word size of 6 for all integration sites, except for the Direct Repeat integration site, which was identified using the blastn-short option. Only BLAST hits covering 70% of the sequence with a 70% identity were used for this analysis. This analysis was validated using long-read sequences from 12 different isolates.

### Pf core genome assembly

When a single phage was found, the boundary of the core genome was defined as the start of the excisionase on one end and the end of the integrase on the other end.

When multiple phages were found, raw reads were aligned to consensus reference genomes corresponding to the type of Pf phages found in that isolate using Bowtie2. We used ten consensus reference genomes made from phage genomes in Fiedoruk et al. (13), corresponding to all possible combinations of the two PA0723 isoforms and the five different PA0727 integrases. A consensus sequence for each phage in our dataset was then obtained by aligning raw reads to the appropriate reference consensus sequence. Phage sequences from multiple phages with different PA0723 isoforms cannot be resolved and reconstructed from short reads when present in the same isolate due to highly conserved areas between PA0723 and the integrase. We omit these phages from the analysis of Pf pairwise distances presented in this work. This method and these parameters were validated using long read sequencing data for 12 samples from the California patient cohort.

### Multiple sequence alignment and phylogenetic tree

Bacterial sequences were aligned and a phylogenetic tree was constructed from the aligned core genomes with Harvest (64). PA7 (BioSample: SAMN02603435) was used as an outgroup. Pf sequences were aligned with Mafft with --adjustdirection (65) and maximum likelihood phylogenetic trees were constructed with FastTree (66). Pairwise distances were calculated using snp-dists.

### Statistics

All statistics were performed with R. T-tests and Wilcoxon Rank-Sum tests were used to compare the means and medians of normal and non-normal distributions, respectively. Chi-square tests were used to compare differences in proportions between two groups.

### Study approval

*Pa* isolates from respiratory cultures from individuals with CF were identified and banked with patient consent for biobanking under IRB #11197.

### Data availability

Raw reads for *Pa* isolates from the California cohort can be accessed through NCBI SRA under BioProject accession PRJNA1188603.

## Author contributions

JDP, PLB and EBB designed the research study. CM and EBB acquired samples. AG, AK, HM, PSP and EBB performed experiments. JDP, NHH, AKS, PF, RJ and TC analyzed the data. All authors contributed to the writing of the manuscript and the design of the figures.

## Acknowledgments

EBB was supported by NHLBI/NIH K23 (1K23HL169902-01) and the Cystic Fibrosis Foundation (BURGEN23G0 & BURGEN24A0-KB). PLB discloses support from National Institutes of Health grant R01 HL148184-01, National Institutes of Health grant R01 AI12492093, National Institutes of Health grant R01 DC019965, Cystic Fibrosis Foundation grant, and Emerson Collective grant. The contents are those of the authors and do not necessarily represent the view of the funding agencies.

