## Supplemental Figures for "*Pseudomonas* superinfection drives Pf phage transmission within airway infections in patients with cystic fibrosis"

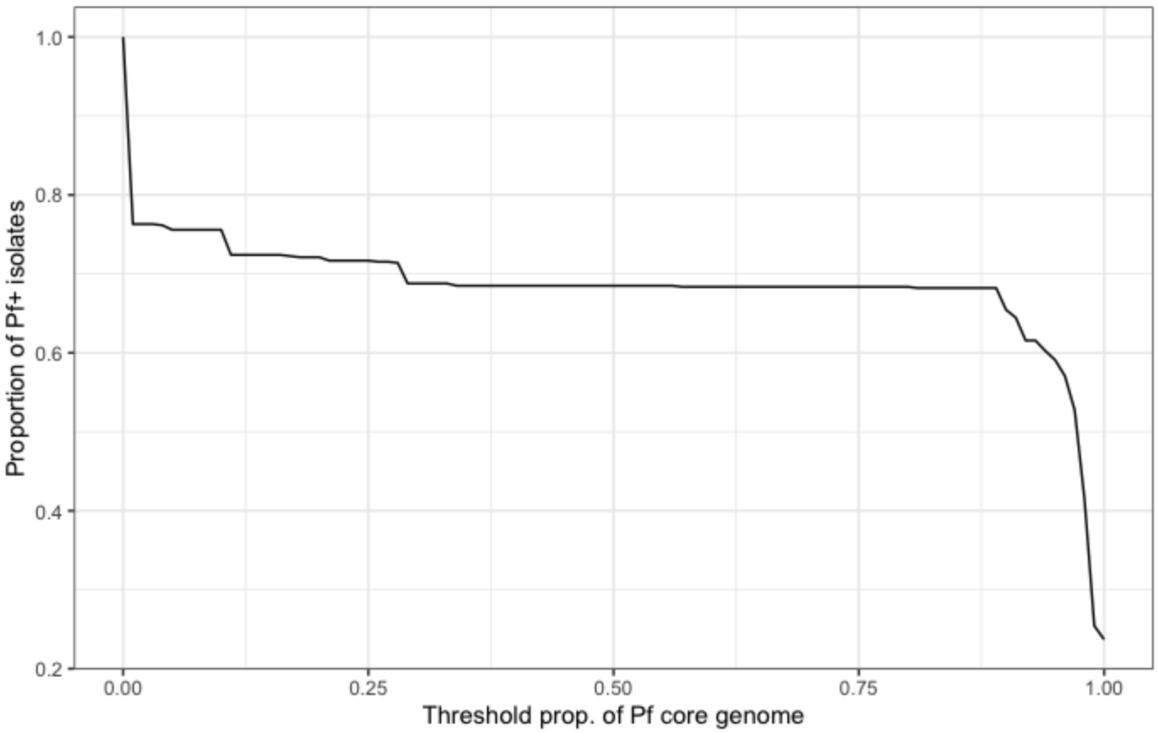


**Figure S1.** Sensitivity of the proportion of Pf+ isolates to core Pf genome threshold required for Pf identification. The core Pf genes considered are PA0718, PA0719, PA0720, PA0721 and PA0727. This analysis included all samples across all three clinical cohorts.

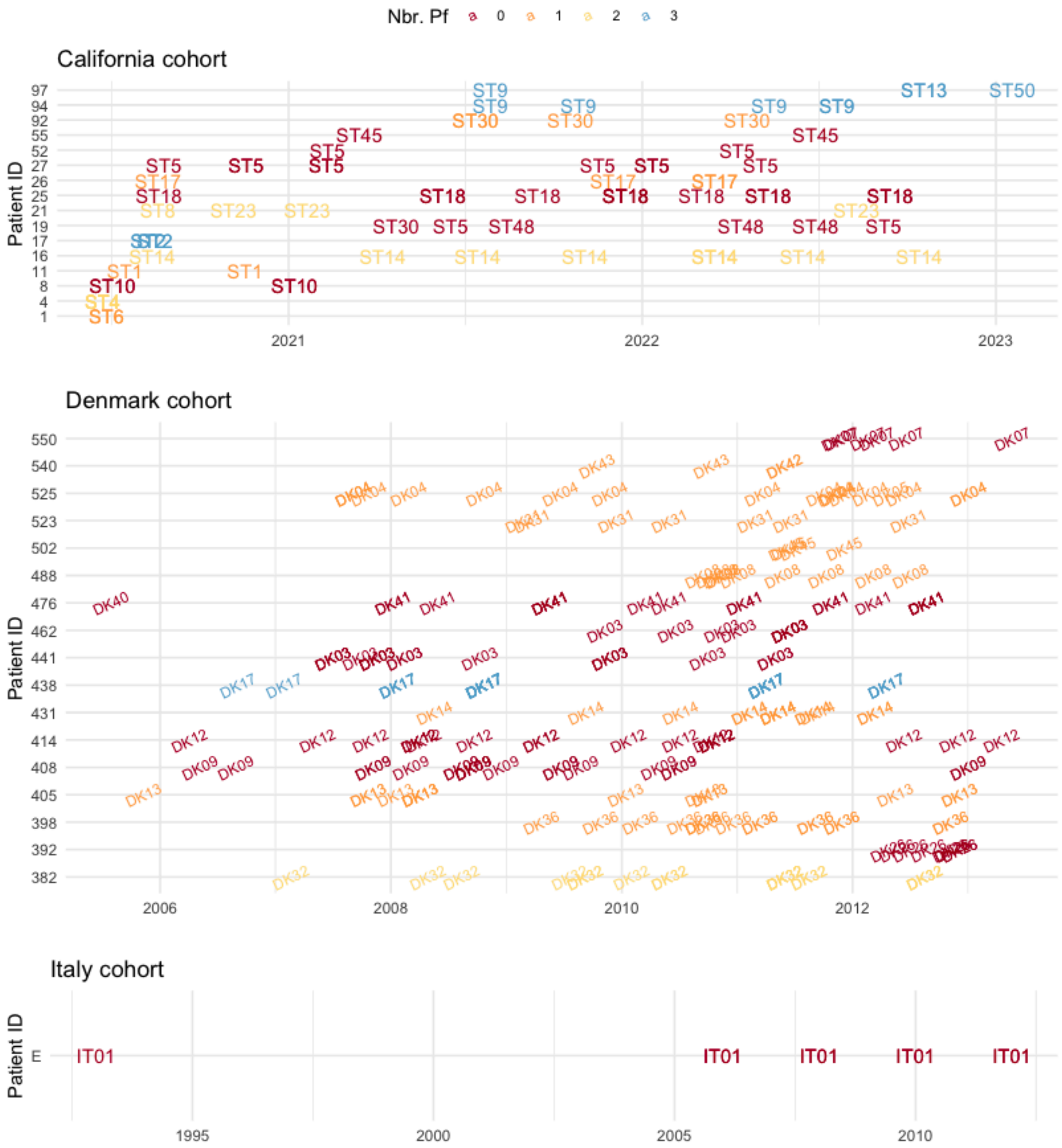


**Figure S2.** Time series of patient samples for patients with isolates with the same number of Pf phages over time. Labels indicate the clone type of each isolate. Colors show the number of Pf prophages in that isolate. Patients with a single sample are not shown.
